# Bacterial clearance enhanced by α2, 3- and α2, 6-sialyllactose via receptor-mediated endocytosis and phagocytosis

**DOI:** 10.1101/414177

**Authors:** Jimin Kim, Yong Jae Kim, Jae Wha Kim

**Author notes:** **Corresponding Author:** Jae Wha Kim, Ph. D. Cell Factory Research Center, Division of Systems Biology and Bioengineering, Korea research Institute of Bioscience and Biotechnology, Daejeon 34141, Republic of Korea. Jimin Kim and Yong Jae Kim contributed equally to this work.

## Abstract

Sialyllactose (SL) is a representative human milk oligosaccharide (HMO) of human breast milk. The roles of SL in infant brain development and immunity have been reported in previous studies. In this study, we identified the impact of SL on innate immunity. Our results showed that the administration of SL had significant efficacy on bacterial clearance in *Pseudomonas aeruginosa K*-infected mice. We also examined the role of SL in the human THP-1 macrophage-like cell line. SL effectively promoted receptor-mediated endocytosis and phagocytosis. Furthermore, SL accelerated the recruitment of Rac1 to the cell membrane, leading to the generation of reactive oxygen species for the elimination of phagocytosed bacteria. Our findings provide a new perspective on the role of SL in breast milk and suggest its potential application as a therapeutic agent to treat bacterial and viral infections.

## Introduction

Human breast milk contains considerable amounts of oligosaccharides, defined as human milk oligosaccharides (HMOs). HMOs are present at levels ranging from 15–23 g/L in colostrum and 8–12 g/L in transitional and mature milk (1); the quantity of oligosaccharides present in human breast milk is about 10 to 100 times higher than that in milk from other mammals (2, 3). Because HMO-rich human milk provides the best nutrition for infants, as well as elicits immunity-enhancing and prebiotic effects, the market for the production of nutritional infant foods and healthy functional foods related to HMOs is steadily growing. HMOs comprise five types of monosaccharides: D-glucose, D-galactose, L-fucose, N-acetylglucosamine, and sialic acid (N-acetylneuraminic acid); these monosaccharides compose complex and various HMOs (4). The structures and types of HMOs are very diverse, with about 200 oligosaccharides found in breast milk. Among them, sialyllactose (SL), which is composed of sialic acid and lactose, is classified into α2, 3-SL and α2, 6-SL, depending on the position at which the sialic acid binds lactose (5).SL is a major source of sialic acid, a component of the gangliosides present on neuronal surfaces (6, 7), and is essential for brain function and cognitive development (8, 9). SL is also known to inhibit the adhesion of pathogens to the intestinal epithelium, thus preventing pathogenic infections (10). SL is thought to be involved in the regulation of the immune system, but specific experimental results are lacking. We therefore aimed to confirm the efficacy of SL in innate immunity.

Innate immunity is the primary line of defense against infective organisms. Macrophages, phagocytes such as neutrophils, and skin barriers all play important roles in innate immunity (11). Macrophages are immune cells that remove infectious pathogens through phagocytosis (12). Phagocytosis, a specific type of endocytosis, is the process by which phagocytes ingest foreign substances and remove them (13). Phagosomes, which are formed via phagocytosis, combine with lysosomes, which contain hydrolytic enzymes and reactive oxygen species (ROS), to form phagolysosomes, resulting in the degradation and elimination of pathogens (14). Additionally, after phagocytosis occurs and the toll-like receptor 4 (TLR4) recognizes the bacteria (15, 16), two different signaling pathways are activated: One is the myeloid differentiation primary response 88 (MyD88)-dependent pathway from the TLR4 in the membrane, and the other is the endosomal toll/interleukin-1 receptor (TIR) domain-containing adaptor-inducing interferon β (TRIF)-dependent signal (17). These two signals commonly activate nuclear factor-kappa B (NF-κB) in macrophages to promote the secretion of the C-X-C motif chemokine ligand 8 (CXCL8) (18, 19). Secretion of CXCL8 appears to be evidence of macrophage activation and can recruit other phagocytes like neutrophils to further assist in bacterial clearance (20).

*Pseudomonas aeruginosa K* (PAK), a Gram-negative bacteria, can cause a variety of infectious diseases, such as urinary tract infections (21), gastrointestinal infections (22), skin and soft tissue infections (23), and respiratory infections (24, 25). Respiratory infections caused by PAK can result in pneumonia, which can cause high mortality (26). Therefore, we investigated PAK-induced pneumonia models both *in vivo* and *in vitro*, and determined the effect of SL on disease outcome. We found that SL enhances bacterial clearance via the endocytic activity of macrophages, which suggests that SL might be a novel therapeutic agent for the treatment of pathogen-induced inflammation.

## Results

### Both α2, 3- and α2, 6-SL enhance bacterial clearance in PAK-infected mice

A diagram of the PAK-induced pneumonia mouse model and experimental design are shown in Figure 1A. SL was administered to mice via oral inoculation. Mice were inoculated with PAK via intranasal inoculation, and the amount of PAK in the bronchoalveolar lavage fluid (BALF) was measured, as described above. The clearance of invading PAK was significantly enhanced in both the α2, 3- and α2, 6-SL-treated groups (Fig. 1B).

**Figure 1.**
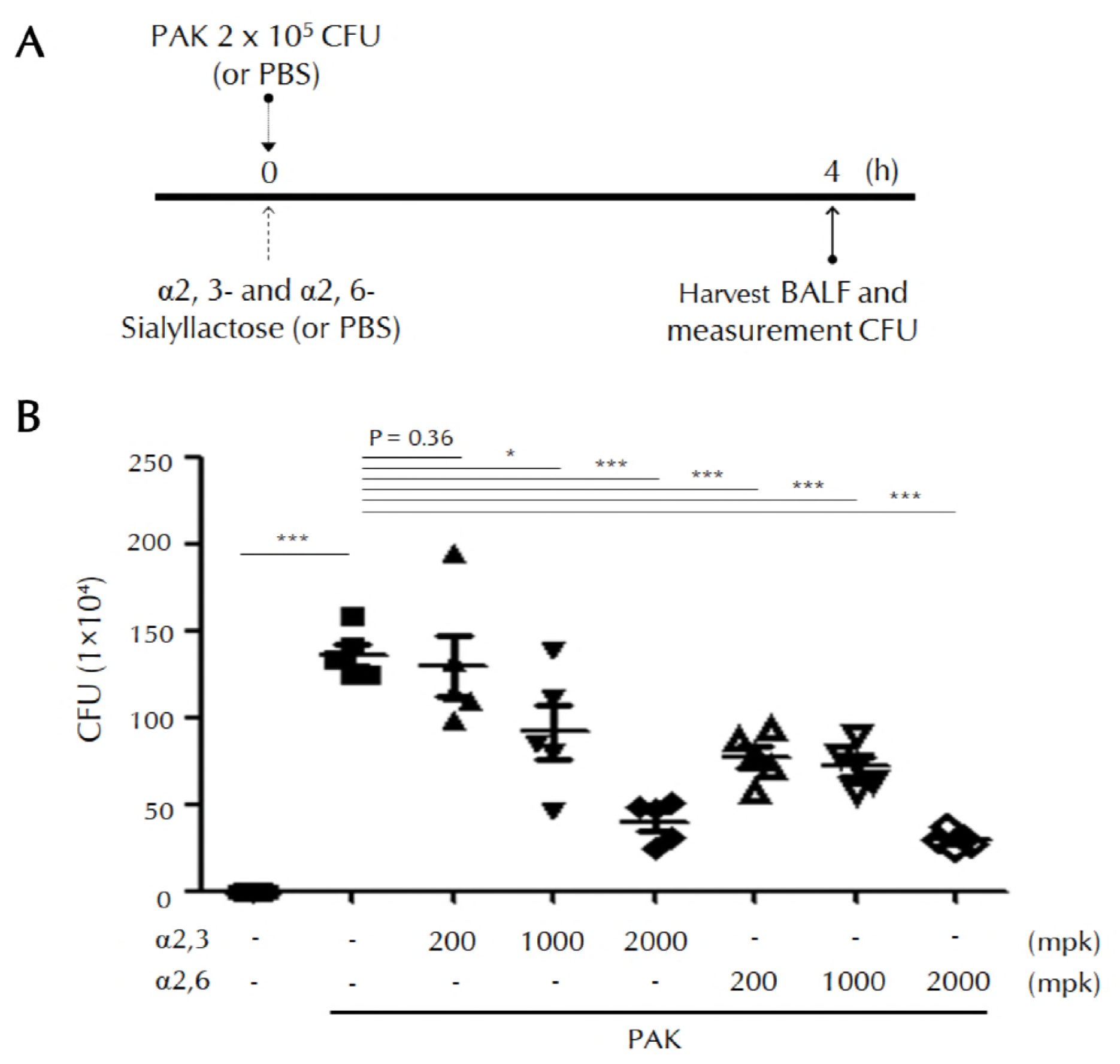
Effects of SL on bacterial clearance in PAK-infected mice. (A) Diagram of the experimental design of a pneumonia mice model. (B) PAK (2 × 10^5^CFU) was administered to mice by intranasal inoculation, and both α2,3- and α2,6-SL (200, 1,000, and 2,000 mg/kg) were administered orally. After PAK infection for 4 h, we harvested the BALF and measured the amount of PAK (CFU). Bars represent the mean ± SD. Control group (PBS); PAK group (PAK only); α2,3-SL-treated group (PAK and α2, 3-SL); α2, 6-SL-treated group (PAK and α2, 6-SL). Five mice were used per group. \**P* < 0.05, \*\**P* < 0.01, \*\*\**P* < 0.005. Abbreviations: PAK, *Pseudomonas aeruginosa K*; CFU, colony forming unit; PBS, phosphate-buffered saline; BALF, bronchoalveolar lavage fluid; α2,3, α2,3-sialyllactose; α2,6, α2,6-sialyllactose; mpk, milligrams per kilogram.

### Both α2, 3- and α2, 6-SL promote phagocytosis of PAK and TLR4 endocytosis in THP-1 cells

Prior to *in vitro* experiments, we examined the cell cytotoxicity of SL by treating the concentration from 0.1 to a maximum of 300 μM (Fig. 2A). Both α2, 3- and α2, 6-SL showed no toxicity in THP-1 cells at all concentrations. Figure 2B shows the experimental design for measuring *in vitro* phagocytic activity. The amount of phagocytosed PAK was significantly increased in SL-treated cells after 1 h. A dramatic decrease in the amount of intracellular bacteria was also detected in SL-treated cells after 2 h (Fig. 2C). These results suggest that internalized PAK might be readily degraded after 1 h. We then investigated TLR4 endocytosis using lipopolysaccharide (LPS) present in the outer membrane of PAK by confocal microscopy. LPS-loaded TLR4 in the cell membrane was internalized within 1 h. In the SL-treated groups, internalization of LPS/TLR4 was observed within 30 min. Both α2, 3- and α2, 6-SL significantly promoted the internalization of TLR4 (Fig. 2D).

**Figure 2.**
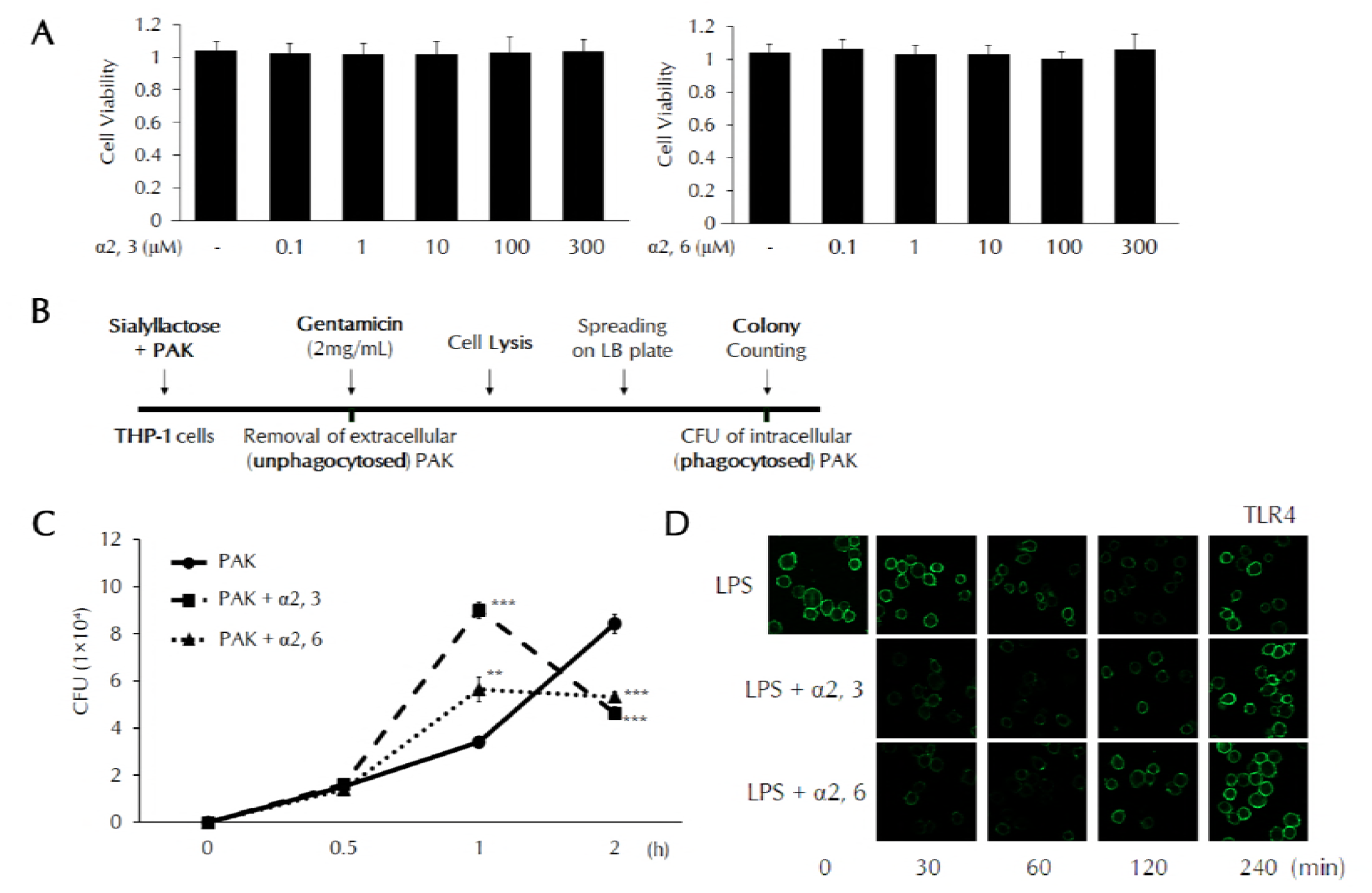
Phagocytosis and TLR4 endocytosis in SL-treated THP-1 cells. (A) Cell cytotoxicity of SL was analyzed by WST-1 assay. (B) Diagram of the measurement of *in vitro* phagocytic activity. (C) THP-1 cells were pretreated with α2, 3- or α2, 6-SL (10 μM) for 3 h, and then treated with PAK at a multiplicity of infection (MOI) of 50 for the indicated times. After PAK infection,we measured the amount of PAK in THP-1 cells. \*\**P* < 0.01 and \*\*\**P* < 0.005 compared to the PAK group. (D) THP-1 cells were pretreated with α2, 3- or α2, 6-SL (10 μM) for 3 h, and then treated with LPS (0.1 μg/mL) for the indicated times. The cells were fixed and incubated with anti-TLR4 antibodies. TLR4 expression in the cell membrane was observed by fluorescence confocal microscopy. Abbreviations: PAK, *Pseudomonas aeruginosa K*; LB, lysogeny broth; CFU, colony forming unit; α2,3, α2,3-sialyllactose; α2,6, α2,6-sialyllactose; LPS, lipopolysaccharide; TLR4,toll like receptor 4.

### Both α2, 3- and α2, 6-SL increase ROS generation and bacterial clearance

Intracellular ROS levels were measured by flow cytometry using DCFH-DA. In the LPS-treated group, ROS levels were maximal after 2 h. Maximal levels of ROS were found after 1 h in SL-treated groups. The return to control levels of ROS was accelerated in SL-treated cells. Mean fluorescence intensity (MFI) values increased more rapidly in the SL-treated group compared to LPS only-treated group, and promptly returned to normal after the elimination of bacteria (Fig.3A). We further analyzed the assembly of Rac1 on the cell membrane. Rac1 is a component of NOX (NADPH oxidase), which generates ROS to remove invading bacteria. During the internalizing of PAK, increased Rac1 assembly was observed at the membrane, which was shown by western blotting and confocal imaging. Membrane-located Rac1 levels increased, whereas cytosolic Rac1 levels decreased, as determined by western blotting analysis, suggesting that Rac1 is translocated to the membrane from the cytosol during bacterial phagocytosis (Fig. 3B). The assembly of Rac1 on the membrane was accelerated after SL treatment. These results were also confirmed by the results of the immunofluorescence assay (Fig. 3C). To evaluate Rac1 localization to the membrane, permeabilization with antibodies was not performed. We then examined intracellular lysosomal activity using the Lyso-tracker dye during phagocytosis (Fig. 3D). Extensive lysosomal activity was found 2 h after TLR4 stimulation by LPS. This activity was observed at 1 h and was significantly enhanced by SL treatment. Our data suggest that SLs play a role in the early eradication of invading bacteria and stimulate a rapid return to homeostasis.

**Figure 3.**
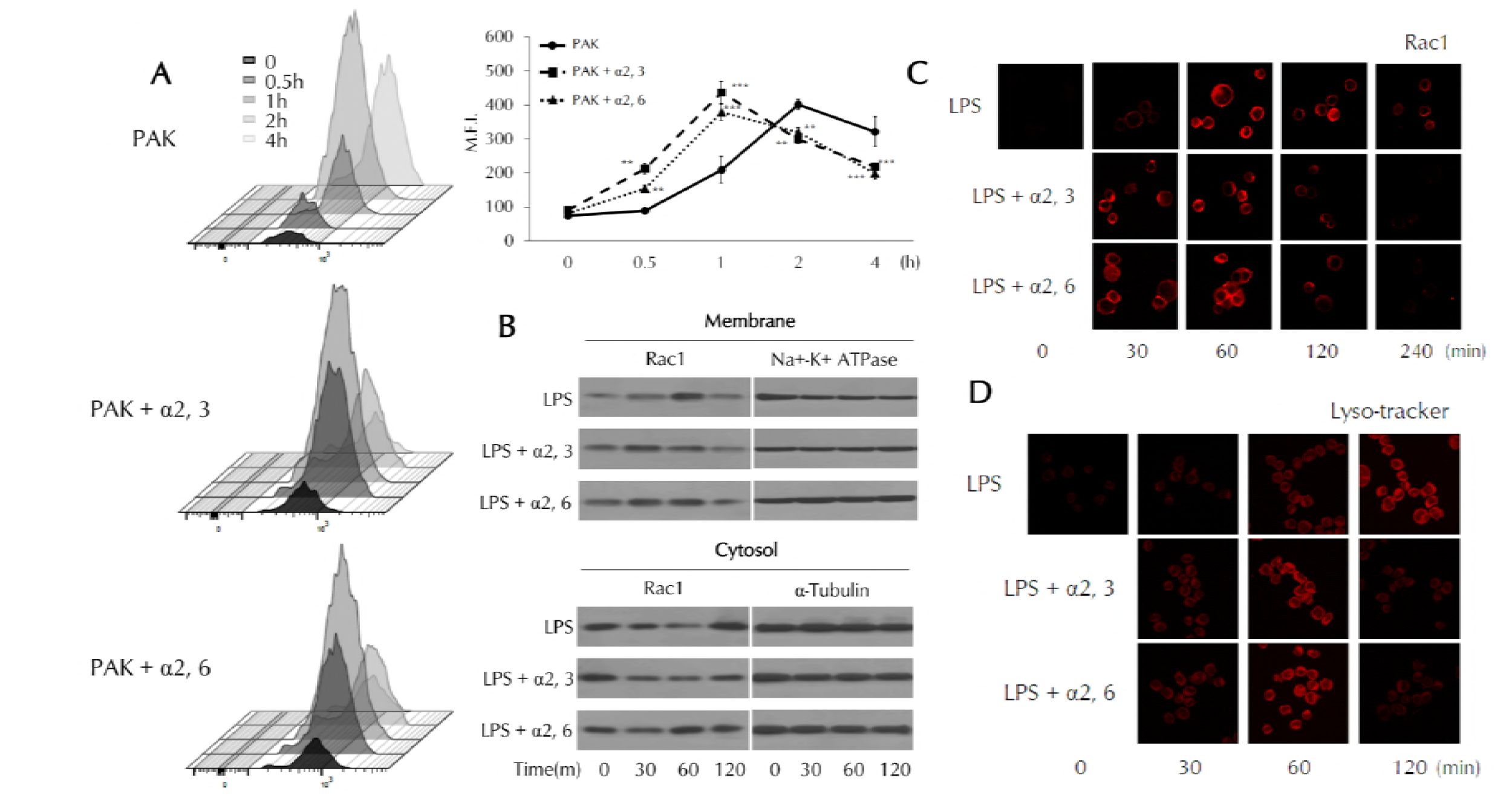
ROS generation and Rac1 localization in SL-treated THP-1 cells. (A) THP-1 cells were pretreated with α2, 3- or α2, 6-SL (10 μM) for 3 h, and then treated with PAK (MOI of 50) for the indicated times. ROS generation and MFI were analyzed by flow cytometry using DCFH-DA. \*\**P* < 0.01 and \*\*\**P* < 0.005 compared to the PAK group. (B) THP-1 cells were pretreated with α2, 3- or α2, 6-SL (10 μM) for 3 h, and then treated with LPS (0.1 μg/mL) for the indicated times. Protein levels of Rac1 in both the membrane and cytosolic fractions were analyzed by western blotting. (C) THP-1 cells were pretreated with α2, 3- or α2, 6-SL (10 μM) for 3 h, and then treated with LPS (0.1 μg/mL) for the indicated times. The cells were fixed and incubated with either anti-Rac1 antibodies or (D) Lyso-tracker dye. Rac1 localization to the cell membrane and lysosomal activity were observed by fluorescence confocal microscopy. Abbreviations: PAK, *Pseudomonas aeruginosa K*; MFI, mean fluorescence intensity; α2,3, α2,3-sialyllactose; α2,6,α2,6-sialyllactose; LPS, lipopolysaccharide.

### The roles of α2, 3- and α2, 6-SL in bacterial clearance are confirmed by CXCL8 expression

CXCL8 transcripts were detected after 15 min in SL-treated cells, but after 60 min in LPS only-treated cells (Fig. 4A). These data indicate that CXCL8 transcriptional activity was increased in SL-treated cells. The secretion of CXCL8 was effectively induced by PAK; SL further increased the secretion of CXCL8 in a dose-dependent manner (Fig. 4B). CXCL8 expression is known to be controlled by NF-κB. IκB degradation, which leads to NF-κB activation, can be detected by western blotting analysis. Increased IκB degradation after SL addition was detected at 20 min (Fig. 4C). Addition of the NF-κB inhibitor, BAY11-7082, completely abolished the increased CXCL8 expression in response to PAK infection and the effect by SL which was verified by previous results (Fig. 4D). We also examined the CXCL8 secretion in order to determine if SL alone was effective (Fig. 4E). Both α2, 3- and α2, 6-SL did not show any increase effect of the cytokine expression only with the SL alone treatment.

**Figure 4.**
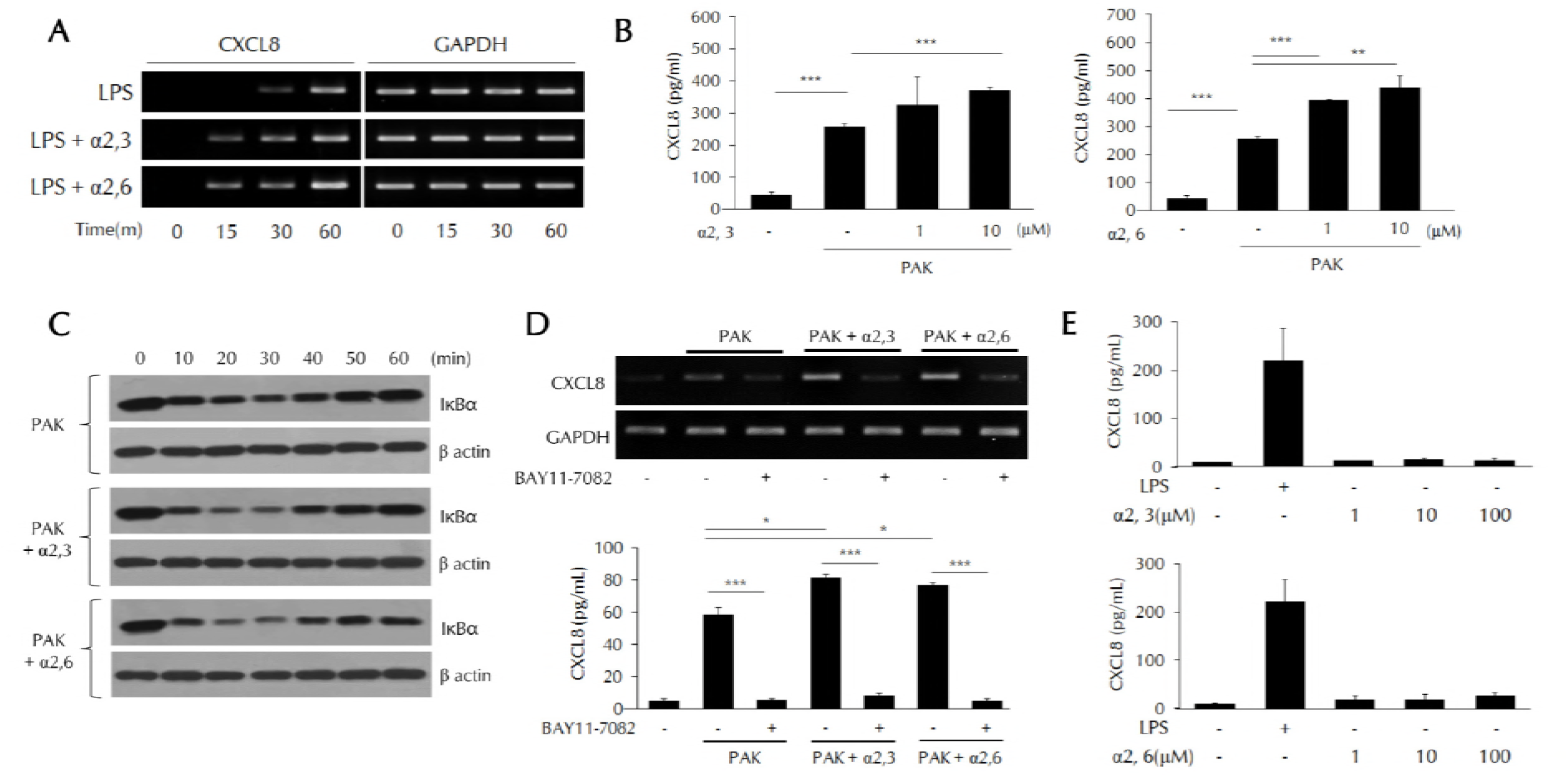
CXCL8 expression via NF-κB activation in SL-treated THP-1 cells. THP-1 cells were pretreated with α2, 3- or α2, 6-SL (1, 10 μM) for 3 h, and then treated with LPS (0.1 μg/mL) or PAK (MOI of 50). (A) CXCL8 mRNA levels were analyzed by RT-PCR. (B, E) CXCL8 secretion was analyzed by ELISA. (C) IκBα degradation was analyzed by western blotting. (D) THP-1 cells were pretreated with the NF-κB inhibitor BAY11-7082 (10 μM), and then treated with α2, 3- or α2, 6-SL (1, 10 μM) and PAK (MOI of 50) as previously described. CXCL8 expression was analyzed by RT-PCR or ELISA. \**P* < 0.05, \*\**P* < 0.01, \*\*\**P* < 0.005. Abbreviations: CXCL8,C-X-C motif chemokine ligand 8; GAPDH, glyceraldehyde 3-phosphate dehydrogenase; α2,3,α2,3-sialyllactose; α2,6, α2,6-sialyllactose; LPS, lipopolysaccharide; PAK, *Pseudomonas aeruginosa K*.

### PAK-induced NF-κB activation by α2, 3- and α2, 6-SLs is dependent on endosome-related signaling

PAK-bound TLR4 at the cell surface activates MyD88 adaptor-related signaling and TLR4 transmits endosome-dependent signals via the activation of TRIF adaptor molecules (27). To identify the role of SL in accelerated endocytosis, we studied the role of SL in these endosome-dependent signals (Fig. 5A). In cells transfected with siRNAs of the endocytosis-related protein clathrin, the acceleration of IκB degradation induced by SL disappeared (Fig. 5B). The increase in CXCL8 expression induced by SL was also not evident. In MyD88 siRNA-transfected cells, CXCL8 secretion was slightly decreased, but the effect by SL was still visible (Fig. 5C). Meanwhile, in TRIF siRNA-transfected cells, CXCL8 secretion was also slightly decreased, but SL treatment had no further effect, indicating that the effects elicited by SL are dependent on TRIF (Fig. 5C). In Figure 2C, TLR4 was internalized by LPS after 60 min; TLR4 returned to the surface by 240 min. LPS/TLR4 internalization and return to the membrane occurred at 30 and 120 min,respectively, in SL-treated cells. TRIF was also similarly detected in cytosolic fractions (Fig. 5D) During endocytosis, TLR4-recruited TRIF was detected in the cytosol. The rapid fluctuations of cytosolic TRIF levels were observed in the SL-treated group. These results indicate that SL promotes the endocytosis of the TLR4 complex and recruitment of TRIF, which results in NF-κB activation and CXCL8 secretion.

**Figure 5.**
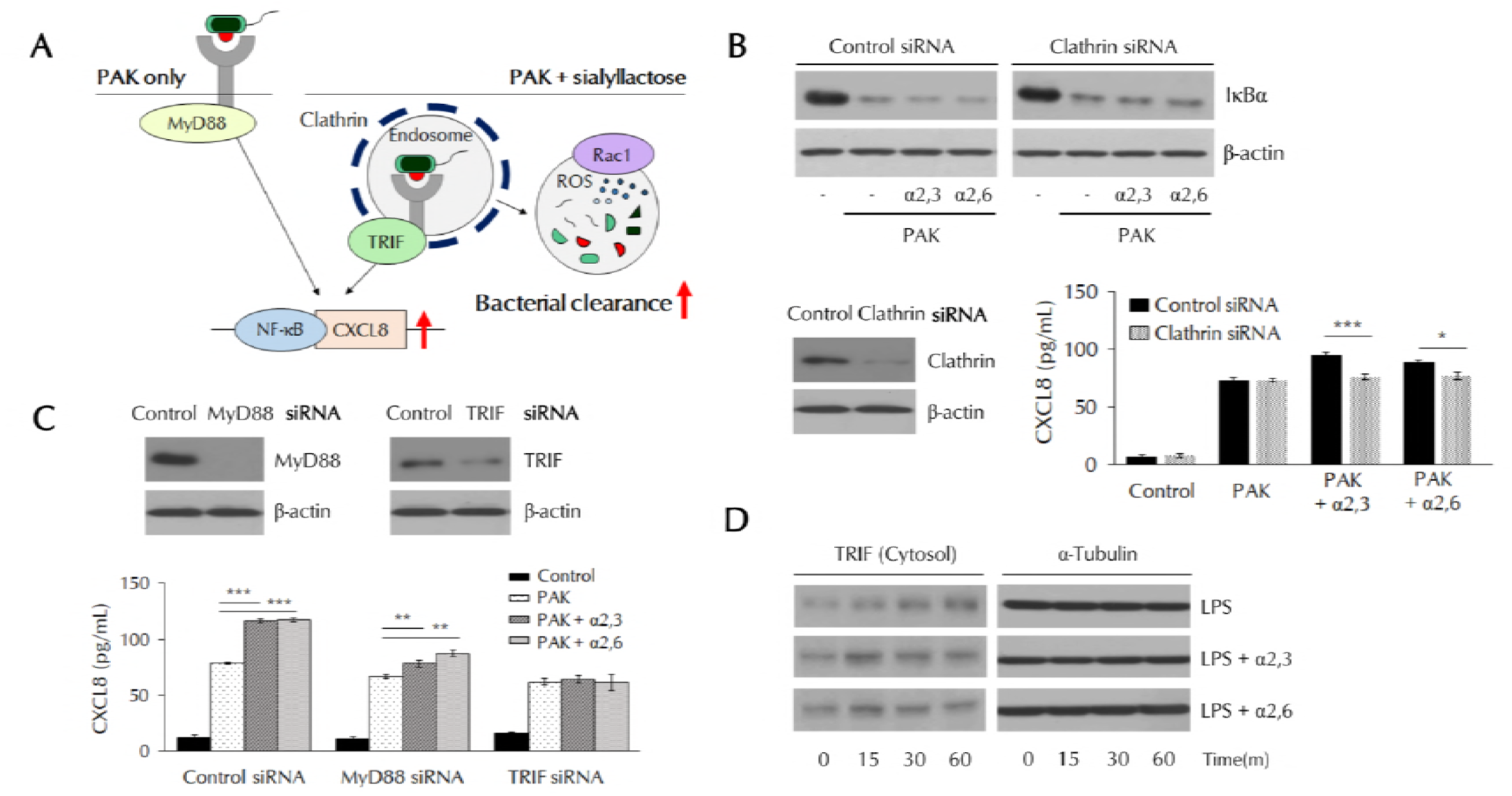
Clathrin-TRIF-dependent NF-κB activation in SL-treated THP-1 cells. (A) Schematic of the experimental design. (B) THP-1 cells were transfected with clathrin siRNA,pretreated with α2, 3- or α2, 6-SL (10 μM) for 3 h, and then treated with PAK (MOI of 50). IκBαdegradation was analyzed by western blotting, and CXCL8 secretion was analyzed by ELISA. (C)THP-1 cells were transfected with MyD88 or TRIF siRNA, pretreated with α2, 3- or α2, 6-SL (10μM) for 3 h, and then treated with PAK (MOI of 50). CXCL8 secretion was analyzed by ELISA.\**P* < 0.05, \*\**P* < 0.01, \*\*\**P* < 0.005. (D) Protein levels of TRIF in cytosolic fractions were analyzed by western blotting. Abbreviations: PAK, *Pseudomonas aeruginosa K*; Myd88, myeloid differentiation primary response 88; TRIF, toll/interleukin-1 receptor (TIR) domain-containing adaptor-inducing interferon; TLR4, toll like receptor 4; NF-κB, nuclear factor-kappa B; CXCL8,C-X-C motif chemokine ligand 8; ROS, reactive oxygen species; α2,3, α2,3-sialyllactose;α2,6, α2,6-sialyllactose.

## Discussion

Downstream signaling by activated TLR4 can occur via two distinct pathways: MyD88-and TRIF-dependent signaling pathways (28, 29). MyD88 is an adapter protein that activates NF-κB in response to almost all TLRs (30, 31). TRIF is another adapter that responds to TLRs and induces the production of interferon beta via the activation of interferon regulatory factor 3 (32). TRIF-mediated signaling occurs at the endosomal membrane after the internalization of TLRs (27). It has been reported that TRIF-dependent signaling plays an important role in LPS-induced macrophage activation (33). We found that SL depends on TRIF signaling in PAK-induced macrophage activation, which supports to this assertion. Many studies insist that NF-κB activity is due mainly to MyD88-dependent signaling (34, 35), but our findings show that NF-κB is also activated by TRIF signaling, which significantly enhances the expression of CXCL8. The secretion of CXCL8, which was increased by the treatment of SL, would invoke the recruitment of other phagocytes, like neutrophils, to provide additional support for bacterial clearance.

Sialic acid can bind to the bacterial surface and also to the host cell membranes (7). Many studies suggest that competition for these binding sites prevents bacterial infection when explain the effects of sialic acid (36, 37). However, if it is merely the role of sialic acid that the bacteria could not attach to host cells, the bacteria that are not eliminated will continue to multiply and this will be a problem for the immune system. Our results showed that the SL treatment significantly increased the bacterial clearance and promoted the phagocytosis of macrophages. This suggests that SL, which can bind both to bacteria and host cells, may have helped macrophages capture the bacteria. Some studies have reported that SL rather supports the adherence of bacteria (38), and that the presence of sialic acid binding receptors in macrophages kills the bacteria by promoting phagocytosis (39). This could be an important clue that not only does SL contained in breast milk play a role in preventing bacterial infection, but also helps to effectively remove the bacteria in the body.

The sustained activation of immune cells can lead to untoward effects, such as the induction of an inflammatory response and the initiation of chronic disease (40, 41). However, to combat pathogenic bacteria, the stimulation of immune cells is essential and should not be inhibited unconditionally. In addition to the activation of immune cells, the maintenance of homeostasis by removal of activated phagocytes is critical. We found that SLs significantly promoted the phagocytosis of bacteria, TLR4 internalization, intracellular ROS generation, lysosomal activity, and NF-κB signaling activation. However, it was also confirmed that all of these activities returned to the control state again within a short period of time. Our results show that the action of SL not only promotes the activation of macrophages but returns the immune system to its original state. This process does not induce a chronic inflammatory response but promotes the activation of immune cells and offers protection from harmful pathogens.

Sialic acid-binding immunoglobulin lectins (siglecs) are cell surface proteins that bind glycan-containing sialic acid and are related to cell adhesion and cell signaling (42, 43). Siglecs are present on the surfaces of immune cells and contain immunoreceptor tyrosine-based inhibitory motif (ITIM)-containing cytoplasmic regions that downregulate receptor-mediated cell signaling and inhibit immune cell activation (44, 45). Siglecs are also known to be involved in bacterial clearance, recognizing and binding the sialic acid on bacterial surfaces or to specific receptors and then removing the pathogens (46, 47). We thought the internalization of TLRs would be affected by the interaction between the receptor and siglecs. To determine whether PAK-induced receptor internalization was promoted in siglec-silenced cells, we attempted to knock down siglecs using siRNAs (Data were not shown) for siglec 5, 7, and 9, which are expressed in macrophages and recognize SL (48-50). The results showed that the amount of internalized PAK was enhanced in all siglec-silenced cells. In addition, intracellular ROS generation, IκB degradation and CXCL8 expression were also enhanced in siglec-silenced cells. Our results somewhat differ from those of other studies (42, 43) but suggest another role for siglecs.

At present, there are few studies using free state of SL. However, there is a large amount of free HMOs in breast milk, suggesting the need to investigate the effects of free SL. In this regard, our results provide a new perspective on the action of SL in breast milk and has sufficient research value. In conclusion, SL can be developed and used as a therapeutic agent to treat bacterial and viral infections, including PAK-induced pneumonia.

## Materials and Methods

### Chemicals and reagents

Both α2, 3- and α2, 6-SL were from Genechem Inc. (Daejeon, Republic of Korea) and dissolved in distilled water for all experiments.

### Animal

Six-week-old male BALB/c mice were purchased from KOATECH Corporation (Gyeonggi-do, Republic of Korea). Mice were housed in a specific pathogen-free facility under consistent temperature of 24 °C for 12-hour light/dark cycles. All experimental procedures were approved by the Institutional Animal Care and Use Committee of the Korea Research Institute of Bioscience and Biotechnology and performed in compliance with the National Institutes of Health Guidelines for the care and use of laboratory animals and Korean national laws for animal welfare.

### Bacterial culture and preparation for infection

PAK was cultured in LB (Duchefa Biochemie, Haarlem, Netherlands) or on LB agar plates overnight at 37 °C. Bacterial cells were harvested by centrifugation at 13,000 × *g* for 2 min after an overnight culture. The bacterial pellet was resuspended to 2 × 10^5^ colony-forming units (CFU) in 20 μL phosphate-buffered saline (PBS), as determined by optical density and serial dilution on LB agar plates.

### PAK-infected mouse model

PAK was administered to mice by intranasal inoculation; SL (200, 1,000, and 2,000 mg/kg) was administered to mice by oral inoculation. The mice were divided into four groups (five mice per group): control group, PAK-only infected group, α2, 3-SL-treated group, and α2, 6-SL-treated group. Samples of BALF were collected from PAK-infected mice 4 h after infection, and the number of viable bacteria was determined by counting colonies formed on the plate. Harvested BALF was serially diluted to 1:1,000–1:10,000 with PBS. Diluted samples were plated on LB agar and incubated at 37 °C overnight to determine the CFU.

### Cell culture

THP-1 cells (a human leukemia monocyte line) were cultured in RPMI-1640 medium (Welgene, Gyeongsangbuk-do, Republic of Korea) containing 10% fetal bovine serum (Tissue Culture Biologicals, CA, USA), 50 μM β-mercaptoethanol, 100 units/mL penicillin, and 100 μg/mL streptomycin (Antibiotic-Antimycotic Solution, Welgene). The cells were grown in a humidified atmosphere in 5% CO_2_at 37 °C.

### Cell viability assay

THP-1 cells were seeded into 96-well plates and treated with α2, 3- and α2, 6-sialyllactose (0.1 μM, 1 μM, 10 μM, 100 μM and 300 μM) for 24 h. Cell viability was measured by adding 10 μL of D-Plus™ CCK reagent (Dongin LS, Seoul, Republic of Korea). The cells were incubated for 1 h at 37°C, and the optical density (OD) at 450 nm was determined using an EMax Precision Microplate Reader (Molecular Devices, Sunnyvale, CA, USA).

### Measurement of phagocytosed bacteria in THP-1 cells

To remove the bacteria that were not phagocytosed by macrophages, THP-1 cells were treated with 2 mg/mL gentamycin for 30 min and washed three times in ice-cold PBS. The cells were then harvested, lysed in a 0.25% sodium dodecyl sulfate (SDS) solution, and serially diluted to 1:1,000–1:10,000 with PBS. Each sample was then plated on LB agar and incubated at 37°C overnight. The number of bacteria phagocytosed into THP-1 cells was determined by counting the colonies formed on the plate.

### Immunofluorescence assay

THP-1 cells were harvested, washed in PBS, and fixed with paraformaldehyde. With the exception of experiments to investigate lysosomal activity, no additional permeabilization was performed so as to allow the analysis of only the proteins expressed on the cell surface. To prevent nonspecific antibody binding, cells were blocked with 1% bovine serum albumin (BSA) for 1 h at room temperature. After incubation with anti-TLR4 (sc-13593, Santa Cruz Biotechnology, Dallas, TX, USA) or anti-Rac1 (03589, EMD Millipore, Darmstadt, Germany) antibodies (dilution 1:200), or Lyso-tracker (ENZ-51005, Enzo Life Sciences, Farmingdale, NY, USA, dilution 1:1,000), THP-1 cells were stained with Alexa Fluor 488- or 594-conjugated secondary antibodies (Enzo Life Sciences, dilution 1:1,000) and DAPI (Invitrogen, Carlsbad, CA, USA). The stained cells were observed under a Zeiss LSM800 confocal microscope (Carl Zeiss, Jena, Germany).

### ROS analysis

THP-1 cells were incubated with 2 μM DCFH-DA (Invitrogen) for 30 min at 37 °C after washing in PBS for intracellular ROS analysis. The analysis was performed using a BD FACS Verse flow cytometer (BD Biosciences, Franklin Lakes, NJ, USA). The MFI represented intracellular ROS levels.

### Western blotting

THP-1 cells were lysed in RIPA buffer (Lps Solution, Daejeon, Korea) supplemented with protease and phosphatase inhibitors (Thermo Scientific, Waltham, MA, USA). Proteins were separated by electrophoresis through a 12% sodium dodecyl sulfate-polyacrylamide gel and transferred to a polyvinylidene difluoride membrane (EMD Millipore). The membranes were blocked with 5% BSA for 1 h and incubated with primary antibodies (dilution 1:1,000) to Rac1 (03589, EMD Millipore), IκBα (#4814, Cell Signaling Technology, Danvers, MA, USA), TRIF (4596S, Cell Signaling Technology), and β-actin (sc-1616, Santa Cruz Biotechnology). After three washes with PBS containing Tween 20 (PBST), membranes were incubated with horseradish peroxidase (HRP)-conjugated second antibodies (Enzo Life Sciences, dilution 1:5,000) for 1 h at room temperature. Protein bands were detected using the ECL reagent (Thermo Scientific) and visualized on film.

### Enzyme-linked immunosorbent assay (ELISA)

Ninety-six-well microtiter plates were coated with anti-CXCL8 antibodies (BD Biosciences, dilution 1:250) at 4 °C overnight and then blocked with 2% BSA for 1 h at room temperature, followed by the addition of samples. After incubation for 2 h, plates were washed with PBST, and then anti-CXCL8 antibodies and HRP were added (BD Biosciences, dilution 1:250). The tetramethylbenzidine substrate solution was used for the visualization of reactions,which were terminated by the addition of 2 M sulfuric acid. Secreted CXCL8 levels were measured using an EMax Precision Microplate Reader (Molecular Devices) at 450 nm.

### Transfection

Transfection with siRNAs was performed using the HiPerFect reagent (Qiagen, Hilden, Germany), according to the manufacturer’s protocol. Clathrin siRNA (sc-35067), Myd88 siRNA (sc-35986), and TRIF siRNA (sc-106845) were obtained from Santa Cruz Biotechnology.

### RNA isolation and reverse transcription-polymerase chain reaction (RT-PCR)

Total RNA from THP-1 cells was prepared using the Tri-RNA reagent (Favorgen,Kaohsiung, Taiwan), according to the manufacturer’s instructions. The following primers (Macrogen, Daejeon, Korea) were used in this experiment: CXCL8, forward-5’-AGGGTTGCCA ATGCAATAC-3’; reverse-5’-GTGGATCCTGGCTAGCAGAC-3’. Transcribed DNA was amplified using TOPsimple™ DyeMIX-HOT (Enzynomics, Daejeon, Republic of Korea); products were separated by electrophoresis through 2% agarose gels and visualized under UV light using Printgraph 2M (Atto, Tokyo, Japan).

### Statistical analysis

Data are presented as the mean ± standard deviation (SD). The Student’s *t* test was used to calculate the statistical significance of differences between groups. A *P* < 0.05 was considered statistically significant.

## Acknowledgements

JWK designed the study. JK and YJK researched data. JK and JWK wrote the manuscript. JWK reviewed/edited the manuscript. The authors thank Dr. Sun Young Yoon for her review of the manuscript, and GeneChem Incorporation for providing SL for this study. This work was supported by the Korea Research Institute of Bioscience and Biotechnology (KRIBB) Initiative Research Program (KGM5251813 and KGS9001711).

